# Molecular Correlates of Glycine Receptor Activity in Human β Cells

**DOI:** 10.1101/2024.12.02.626463

**Authors:** Amanda Schukarucha Gomes, Cara E. Ellis, Aliya F. Spigelman, Theodore dos Santos, Jasmine Maghera, Kunimasa Suzuki, Patrick E. MacDonald

**Author notes:** Correspondence to: Patrick MacDonald, Alberta Diabetes Institute, LKS Centre, Rm. 6-126, Edmonton, AB, T6G 2R3, Canada, www.bcell.org.

## Abstract

**Objectives:** Glycine acts in an autocrine positive feedback loop in human β cells through its ionotropic receptors (GlyRs). In type 2 diabetes (T2D), islet GlyR activity is impaired by unknown mechanisms. We sought to investigate if the GlyR dysfunction in T2D is replicated by hyperglycemia *per se*, and to further characterize its action in β cells and islets.

**Methods:** GlyR-mediated currents were measured using whole-cell patch-clamp in human β cells from donors with or without T2D, or after high glucose (15 mM) culture. We also correlated glycine-induced current amplitude with transcript expression levels through patch-seq. The expression of the GlyR α1, α3, and β subunit mRNA splice variants was compared between islets from donors with and without T2D, and after high glucose culture. Insulin secretion from human islets was measured in the presence or absence of the GlyR antagonist strychnine.

**Results:** Although gene expression of GlyRs was decreased in T2D islets, and β cell GlyR-mediated currents were smaller, we found no evidence for a shift in GlyR subunit splicing. Glycine-induced currents are also reduced after 48 hours culture of islets from donors without diabetes in high glucose, where we also find the reduction of the α1 subunit expression, but an increase in the α3 subunit. We discovered that glycine-evoked currents are highly heterogeneous amongst β cells, inversely correlate with donor HbA1c, and are significantly correlated to the expression of 92 different transcripts and gene regulatory networks (GRNs) that include CREB3(+), RREB1(+) and ZNF697(+). Finally, glucose-stimulated insulin secretion is decreased in the presence of the GlyR antagonist strychnine.

**Conclusions:** We demonstrate that glucose can modulate GlyR expression, and that the current decrease in T2D is likely due to the receptor gene expression downregulation, and not a change in transcript splicing. Moreover, we define a previously unknown set of genes and regulons that are correlated to GlyR-mediated currents and could be involved in GlyR downregulation in T2D.

**Highlights:**

- Glycine-evoked currents in β cells are lower in T2D and after high glucose culture
- GlyR gene expression is reduced in T2D, with no shift in splice variant expression
- GlyR inhibition with strychnine decreases insulin secretion
- We define transcripts and gene regulatory networks correlated to glycine-induced currents

## Introduction

Pancreatic β cells are electrically excitable, and insulin secretion is triggered by the firing of action potentials (1,2). The main stimulus for insulin release is glucose, but it is also influenced by other metabolites present in the plasma, such as amino acids and lipids (3,4), and by neurotransmitters that are present in the pancreatic islets (5,6). Those can be secreted by the autonomic innervation present in the pancreas (7), or by the islet cells themselves (8). In type 2 diabetes (T2D), normal neurotransmitter signalling in islets is impaired (9,10), and some neurotransmitters have been studied as possible targets for the treatment of this disease (11,12). Glycine is one of the neurotransmitters present in the islets, where it is secreted by the α and β cells (13,14). Glycine is synthesized from serine in a reaction catalyzed by the enzyme serine hydroxymethyltransferase (SHMT) and then transported from the cytoplasm into secretory vesicles by the vesicular inhibitory amino acid transporter (VIAAT) where it is secreted from the β cell together with insulin (15). The reuptake of glycine from the extracellular space into cells is mediated by the plasma membrane glycine transporters 1 and 2 (GlyT1 and GlyT2) (16). It is also presumed that the circulating glycine from the plasma can reach the pancreatic islets, since glycine ingestion raises plasma insulin in humans (17,18). Interestingly, plasma glycine levels are lower in people who are overweight and with obesity (19–21), and increases after weight loss (22). Glycine is considered a biomarker for T2D, since its plasma concentrations are inversely correlated with the presence and with the risk of developing this disease, even after correcting for BMI (23,24).

The ionotropic glycine receptors (GlyRs) are pentameric ligand-gated ion channels that can be composed of different combinations of its 5 subunits (α1-4 and β), and are permeable to chloride ions (25). These subunits can be alternatively spliced, and both different subunit compositions and different subunit splice variants can result in altered receptor function, or complete inactivity (26–28). Although classically known as an inhibitory neurotransmitter, glycine can be excitatory depending on the chloride balance of the cell (29). This is the case with human β cells, where the chloride equilibrium potential is around -35 mV (30), while the resting membrane potential is -70 mV (2). Our previous work demonstrated that GlyRs are present in human β cells and mediate glycine-evoked currents that can contribute to cell depolarization, increasing insulin secretion (14). β cell calcium responses to GlyR activation are also heterogeneous, either increasing, decreasing, or having no effect on β cell calcium levels (14). The reason behind this heterogeneity and its significance for β cell function is unknown but could be related to variation in GlyR activity or expression. Given that pancreatic β cells are transcriptionally and functionally heterogeneous (31,32), this feature could extend to their neurotransmitter receptors, which has not yet been explored.

Importantly, in islets from donors who lived with T2D, immunostaining for the GlyR α1 subunit is reduced and glycine-evoked currents are decreased (14). The *GLRA1* gene, which encodes the α1 receptor subunit, is also one of the top down-regulated genes in human islets exposed to hyperglycemia (33–36). The mechanisms responsible for those changes and their contribution to impaired insulin secretion in T2D are still unknown. We hypothesized that hyperglycemia *per se* is the specific factor responsible for GlyR dysfunction in T2D. We found that glycine-induced currents are highly heterogeneous among human β cells, even within the same donors. To explore molecules and intracellular pathways involved with GlyR signaling, we sequenced the transcriptome of the β cells after electrophysiologic recording using patch-seq (37), and found a previously unknown set of transcripts that correlate with glycine-evoked current amplitude. We confirmed reduced glycine-evoked currents in T2D β cells, demonstrated a relationship to donor HbA1c, and cultured β cells in high glucose (15 mM) to simulate the chronic exposure to high glucose that happens in diabetes. We found that 2-day high glucose culture reduces glycine-induced currents, indicating that hyperglycemia affects GlyRs. Finally, we investigated if the GlyR dysfunction was mediated by changes in receptor splicing, by quantifying the mRNA expression of all GlyR subunit splice variants in T2D islets and after high glucose culture. We uncovered that GlyR gene expression is overall decreased in T2D and after high glucose culture, but there is no shift towards the expression of non-functional receptor subtypes.

## Methods

### Cell Culture

Human islets were from the Alberta Diabetes Institute (ADI) IsletCore (38), the Clinical Islet Laboratory at the University of Alberta (39), or the Human Pancreas Analysis Program (40) (**Supplementary Table 1**). All research was performed with the approval of the Human Research Ethics Board at the University of Alberta (Pro00013094; Pro00001754) and written informed consent. Islets were handpicked to purity and cultured in DMEM media (Gibco, Waltham, MA, USA) with 10% FBS (Gibco) and 100 U/mL Penicillin/Streptomycin (Gibco) at 37°C with 5% CO_2_. For the electrophysiology experiments, islets were dispersed into single cells using an enzyme-free Hanks’-based cell dissociation buffer (Gibco) and plated into 35 mm culture-treated Petri Dishes (Corning, NY, USA).

### Electrophysiology

Single cells dispersed from human islet donors without diabetes were cultured for 24 or 48 hours in DMEM media with 5.5 mM or 15 mM glucose, while cells from donors with T2D were cultured in the control glucose concentration (5.5 mM). Glycine-mediated currents were measured using an EPC-10 amplifier and the PatchMaster software (HEKA Instruments, Lambrecht/Pfalz, Germany) through the whole-cell patch-clamp technique, at a holding potential of -70 mV. Glycine (300 µM) with and without the GlyR inhibitor strychnine (10 µM) were applied using an SF-77C Perfusion Fast-Step System (Warner Instruments, Hamden, CT, USA). Cells were continuously perfused with an extracellular solution containing: 118 mM NaCl, 20 mM tetraethylammonium-Cl, 5.6 mM KCl, 2.6 mM CaCl_2_, 1.2 mM MgCl_2_, 5 mM HEPES, and 6 mM glucose (pH adjusted to 7.4 with NaOH). Patch-pipettes were pulled from borosilicate glass (Sutter, Novato, CA, USA) with a resistance of 3–6 MΩ when filled with pipette solution containing 130 mM KCl, 1 mM MgCl_2_, 1 mM CaCl_2_, 10 mM EGTA, 5 mM HEPES, and 3 mM MgATP (pH adjusted to 7.2 with KOH). The identity of the cells was later confirmed by insulin (Dako anti-insulin, IR002, Agilent Technologies, 1:5 as primary and Alexa Fluor 488 goat anti-guinea pig, A11073, Invitrogen, 1:200 as secondary) and glucagon (Mouse anti-glucagon, G2654, Sigma, 1:200 as primary and Alexa Fluor 594 goat anti-mouse, A-11032, Invitrogen, 1:200 used as secondary) immunostaining. Analysis was performed using the FitMaster software (HEKA Instruments).

### Patch-Seq

Electrophysiology was performed as described above in human islet cells. Immediately after recording, the patching pipette was replaced by a thin wall borosilicate glass pipette (Sutter), filled with 0.5 µL of lysis buffer (5% Poly-ethylene Glycol 8000, 0.1% Triton X-100, 0.5 u/µL RNAse Inhibitor, 0.5 µM OligodT30VN, dNTPs 0.5 mM/each, Nuclease Free Water), and the cell was collected by applying negative pressure through gentle suction. The cell was then transferred into an 8-strip PCR tube on ice containing 3 µL of lysis buffer, and was stored at -80°C until single-cell RNA sequencing (scRNA-seq) was performed using a modified SmartSeq-3 (41) protocol. Libraries were sequenced with the NextSeq 500 Sequencing System (Illumina). Fastq files were trimmed using TrimGalore version 0.6.5 with auto-detection of adapter sequence, then aligned to the GRCh38 human genome, Ensembl release 104, and counted using STAR version 2.7.11a (42). Decontamination was performed using decontX from the R package celda (version 1.20.0). Cell types were identified using the MapQuery function from the R package Seurat (version 5.1.0), using previously published PatchSeq data as a reference (GSE124742 (37), GSE164875(43), and GSE270484; at PancDB; and at https://explore.data.humancellatlas.org/projects/8559a8ed-5d8c-4fb6-bde8-ab639cebf03c). The pySCENIC (44) workflow was run on decontaminated counts similarly to previously described (45). For Spearman Rank Correlations, transcript expression was normalized to counts per million (CPM) and transformed into Log_2_[CPM+1] values. Cells that had >30% of mitochondrial genes and less than 200 unique transcripts expressed were excluded. Spearman Rank Correlations were performed between current amplitude and expression level for genes detected in at least 50% of cells. Cells with no detectable expression of a given transcript were only excluded from that specific correlation. Single-cell RNAseq data, SCENIC AUC, and electrophysiology parameters were then N-integrated using the R package mixOmics (version 6.28.0) DIABLO workflow (46), comparing beta cells from donors with T2D to beta cells from non-diabetic donors. Transcripts were selected based on Spearman Rank correlations >0.1; all electrophysiology parameters were included; and SCENIC regulons were selected based on the features with the greatest variability.

### qPCR

The RNA expression of the GlyR α1 (*GLRA1*), α3 (*GLRA3*), and β (*GLRB*) subunit splice variants, and of the splicing factors *PTBP1*, *PTBP2*, *NOVA1*, *NOVA2*, *RBFOX1*, *RBFOX2*, and *ELAVL4* genes was measured through the quantitative PCR (qPCR) method (primers in **Supplementary Table 2** and Taqman probes in **Supplementary Table 3**). Fresh islets were used for high glucose treatment, and RNA was extracted after islets from donors without diabetes were cultured in DMEM media (Gibco) in control (5.5 mM) or high (15 mM) glucose for 48 hours. For experiments that quantified gene expression in T2D, RNA was extracted from snap-frozen islet samples of donors with and without T2D from the ADI IsletCore. RNA extraction was performed with the TRIzol reagent (Thermo Fisher) for all samples. To analyze the quality of the extracted RNA, 1µL of each sample obtained was quantified with the Nanodrop 1000 instrument (Thermo Fisher) and the 260/230 and 260/280 values were evaluated. The cDNA was synthesized from 2µg of RNA samples with the One Script Plus cDNA Synthesis Kit (ABM). The qPCR reactions were made using either the Applied Biosystems™ PowerUp™ SYBR™ Green Master Mix (Thermo Fisher) or the Applied Biosystems Taqman Fast Advanced Master Mix (Thermo Fisher) with TaqMan Assays. The gene expression was then quantified by the 2-ΔΔCT method and normalized to the housekeeping gene Cyclophilin A (*PPIA*).

### siRNA Transfection

Human islets dispersed into single cells from donors without diabetes were transfected with *EIF4EBP1* siRNA (Thermo Fisher, siRNA ID s4579) or control siRNA (Accell Non-targeting Control Pool, Horizon Discovery, Cambridge, UK), together with the Fluorescent-labeled siRNA Silencer™ FAM-labeled Negative Control No. 1 siRNA, using Lipofectamine™ RNAiMAX Transfection Reagent (Thermo Fisher). Glycine-evoked currents were recorded through whole-cell patch-clamp after 3 days, as described above, and the identity of the cells was later confirmed by insulin and glucagon immunostaining. RNA was extracted from cells 3 days after transfection using TRIzol (Thermo Fisher), and qPCR was performed to confirm *EIF4EBP1* gene knockdown using TaqMan Assays (Thermo Fisher).

### Dynamic Glucose-Stimulated Insulin Secretion

Insulin secretion of human islets was evaluated by perifusion with a Biorep Perifusion Machine (BioRep, Miami, FL, USA). 25 islets per chamber were perifused at 37°C with a KRB solution containing: 115 mM NaCl, 5 mM KCl, 24 mM NaHCO_3_, 2.5 mM CaCl_2_, 1 mM MgCl_2_, 10 mM HEPES, and 0.1% BSA (pH adjusted to 7.4 with NaOH), with a step-increase in glucose concentration, both in the absence and presence of 10 µM strychnine. After a 30-minute pre-incubation with 2.8 mM glucose, the perifusion was performed according to the following protocol: 10 minutes 2.8 mM glucose, 10 minutes 5.6 mM glucose, 10 minutes 8.4 mM glucose, 10 minutes 11.2 mM glucose, and 16 minutes 2.8 mM glucose. Supernatant samples were collected every 2 minutes. At the end of the experiment, islets were collected in 500 µL of acid ethanol solution (75% ethanol, 23.5% acetic acid, 1.5 % concentrated HCl), and all samples were stored at -20°C until insulin measurement. The secreted insulin and remaining insulin inside the islets (used as the content) were quantified using an ELISA Kit (Alpco Stellux, Salem, NH, USA), and results were calculated as the percentage of secreted insulin in relation to the content.

### Data Analysis

Except for single-cell RNA sequencing analysis, statistical analysis was performed in GraphPad Prism 9 and data is shown as mean ± standard error of the mean (SEM). P values < 0.05 were considered statistically significant. Statistical difference between two groups was determined by paired or unpaired t-test, or non-parametric Mann-Whitney test. Insulin secretion data was analyzed by Two-Way Mixed ANOVA. Specific statistic tests are described in each figure legend.

### Data Availability

Patch-seq data is deposited on the NCBI gene expression omnibus (GEO) with accession number GSE280267. Other data is available upon request and/or available at www.humanislets.com. Code for scRNA-seq analyses is available at https://github.com/caraee/glycine_patchseq.

## Results

### Glycine-evoked currents in β-cells are decreased in T2D and GlyR inhibition lowers glucose-stimulated insulin secretion

First, we recorded glycine-evoked currents to evaluate the activity of the GlyRs in human β cells (**Figure 1A,B**). The glycine-evoked currents measured in β cells from donors with T2D were smaller than those in cells from donors with no diabetes (**Figure 1C**), confirming our previous findings (14). To evaluate the effect of the glycine receptors on insulin secretion, we perifused human islets from donors without diabetes with step-increases in glucose both in the absence and presence of the GlyR inhibitor strychnine (**Figure 1D**). In the presence of 10 µM strychnine, glucose-stimulated insulin secretion was still present, but was decreased compared to controls, indicating that the GlyRs influence insulin secretion (**Figure 1E**).

**Figure 1.**
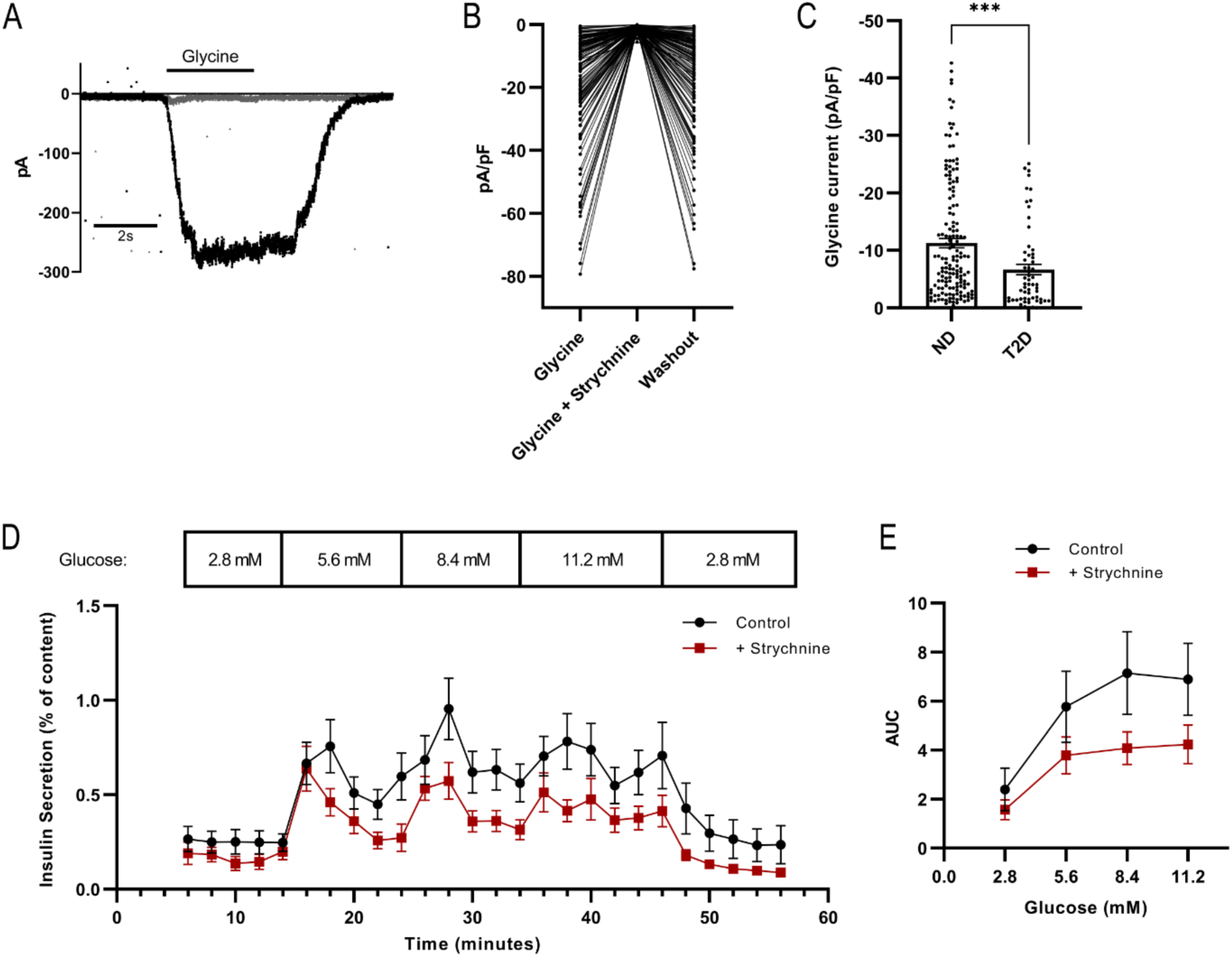
Blockade of heterogenous glycine-evoked currents in human β cells reduces insulin secretion. A: Representative trace recorded (in black) of a glycine (300 µM)-evoked current, and in gray of glycine with the addition of strychnine (10µM). B: Glycine-evoked currents measured in human β cells, followed by I_Gly_ inhibition with 10µM strychnine and washout glycine current (n = 145 cells from 36 donors). C: Currents from donors with (n = 61 cells from 13 donors) and without T2D (ND, n = 160 cells from 43 donors). Data was compared by the Mann-Whitney test, p = 0.0004. D: Percentage of insulin secretion in relation to the content in human islets, in the absence and presence of 10 µM strychnine (n = 6 donors). E: Area under the curve (AUC) calculated for insulin secretion in the absence (23.83 ± 2.43) and in the presence (13.58 ± 1.10) of strychnine (difference calculated by Two-Way Mixed ANOVA, p = 0.035).

### Heterogeneous glycine-evoked currents are associated with different gene transcripts

To further explore the heterogeneity of the GlyRs and uncover transcriptional pathways associated with GlyR signalling and modulation, we measured single-cell mRNA expression of the β cells after current recording with the patch-seq method (**Figure 2A**), and correlated transcript levels with glycine-induced current amplitude. We found that there is a wide variation of glycine-evoked current amplitudes between β cells, even within the same donors (**Figure 2B**). We then applied Spearman Rank Correlations between transcript expression and electrophysiological parameters to the β cells that passed quality control (37,43). Although we did not detect GlyR subunit expression in this assay, we found that glycine-evoked currents are significantly correlated or anticorrelated to the expression of 92 different transcripts (**Supplementary Table 4**). Importantly, these did not correlate significantly with cell size, and correlations were lost in the presence of the inhibitor strychnine and mostly recovered upon washout, indicating true correlation with GlyR activity (**Figure 2C**). Among correlated transcripts are several genes known to influence β cell function, including enzymes (Stearoyl-CoA desaturase - *SCD*), membrane receptors (Adhesion GPCR-G1/GPCR56 – *ADGRG1*), RNA binding proteins (Heterogeneous nuclear ribonucleoprotein K - *HNRNPK*, Cold inducible binding protein - *CIRBP*, Polypyrimidine tract-binding protein 1 - *PTBP1*), translation factors (Translation initiation factor 4E-binding protein 1 - *EIF4EBP1*), and proteins related to exocytosis (Syntaxin-1A *STX1A*, Chromogranin B – *CHGB)* (**Figure 2D**).

**Figure 2.**
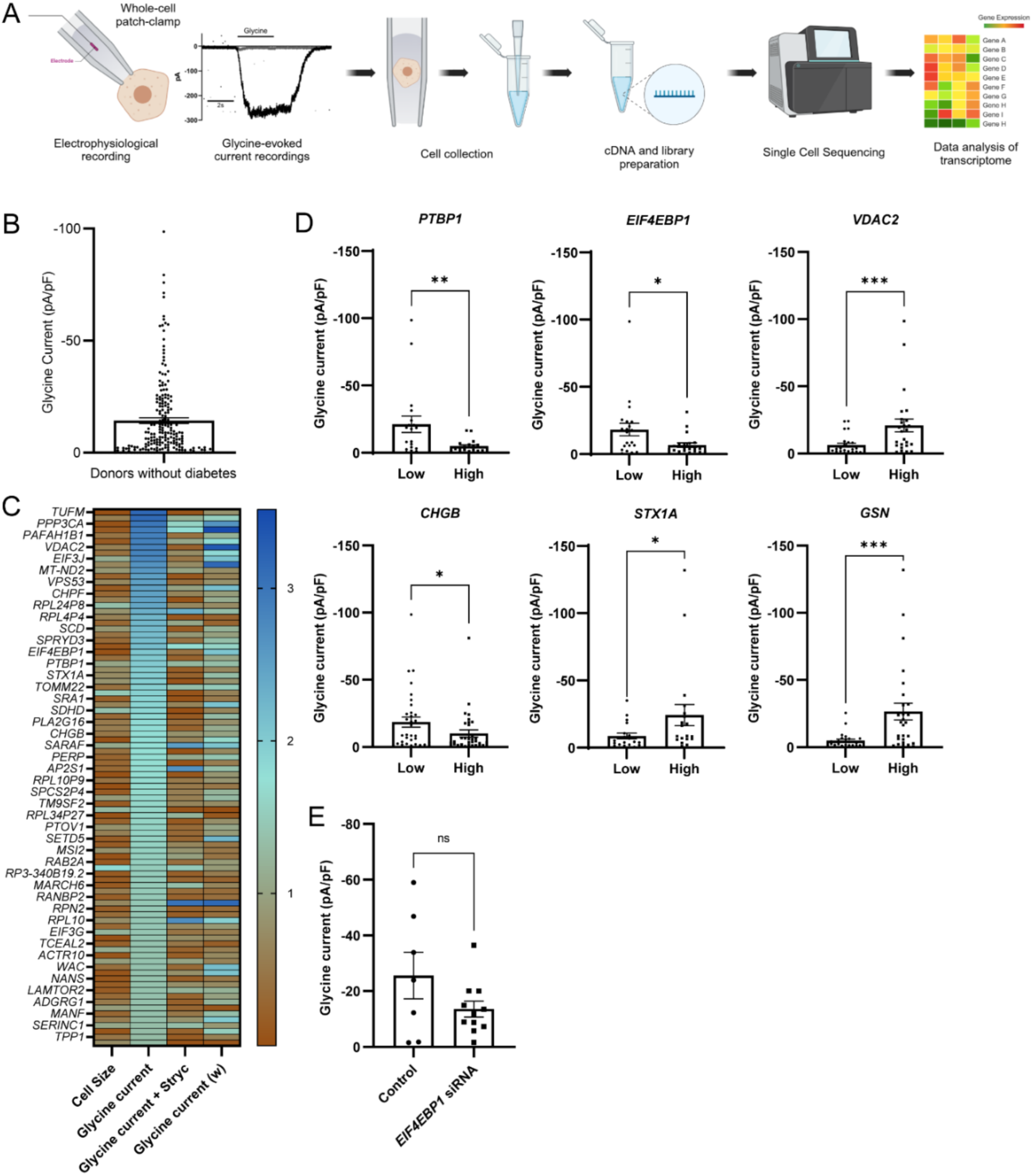
Transcripts correlated to glycine-induced current amplitude in human β cells. A: Schematic of Patch-Seq pipeline. B: Distribution of glycine-evoked current amplitudes in β cells from donors without diabetes (n=176 cells from 43 donors). C: Heatmap of genes significantly correlated to glycine-evoked currents in human β cells. P values for each transcript correlation to cell size, glycine-evoked current, glycine-evoked current in the presence of strychnine, and washout glycine current were transformed into Y = -1*Log(Y). D: Glycine-evoked currents in subpopulation of cells with higher half and lower half expression levels of correlated transcripts. Data was compared by Mann-Whitney test. E: Glycine-evoked currents measured in cells transfected with control siRNA (n=7 cells) or EIF4EBP1 siRNA (n=11 cells from 4 donors). Data was compared by Mann Whitney test, p = 0.3749. * P ≤ 0.05 and ** P ≤ 0.01.

Because the Translation Initiation Factor 4E-binding Protein 1 (*EIF4EBP1*) correlated with glycine-evoked currents, controls translation initiation, and could in theory impact GlyR protein expression, we decided to test if knocking down *EIF4EBP1* in human β cells would result in a change in glycine-evoked current amplitude (**Figure 2E**). However, we found that the average glycine-evoked current is not significantly affected by silencing *EIF4EBP1* expression in human β cells, suggesting that it is not causative of reduced GlyR activity in the T2D β cells.

To further understand the transcriptional differences between the non-diabetic and T2D β cells, we applied SCENIC to infer transcription factor activity and create transcription factor-specific gene regulatory networks, identifying which regulons (the transcription factor and the inferred genes in that pathway) are active in each population (**Figure 3A, Supplementary Table 5**). In agreement with previous results, we observed decreased β cell transcription factor activity with lower Z and regulon specificity scores in T2D β cells, including MAFA(+) and FOXO1(+) (47). We did not observe any regulons that were more specifically active in T2D β cells compared to non-diabetic β cells. We then performed a multiomics analysis combining transcript expression, transcription factor activity, and electrophysiological parameters, with the goal of elucidating the relationship between glycine receptor current amplitude and the overall transcriptional signature of the cells (**Figure 3B, Supplementary Table 6**). The mixOmics N-integration approach selects features from the multi-omics data set to maximize differences between the T2D and non-diabetic β cells within each data type, then can assess how these features vary in the multi-omics context to establish population signatures, similar to biomarkers. Despite selecting transcripts identified based on Spearman Rank Correlation, no transcripts selected by the model were identified as varying with the electrophysiological parameters in this context, reflecting the overall similarity in transcript expression between T2D and non-diabetic β cells and the small sample size. However, we observed a set of regulons that varied similarly to the glycine current amplitude, including OVOL2(+), ZFP3(+), STS1(+), KLF3(+), and ZNF697(+). We also observed a set of regulons that were negatively correlated with glycine current amplitude, including RREB1(+), CREB3(+), and ELF3(+) **(Figure 3C)**, all of which are less active in the T2D β cells compared to non-diabetic β cells. These relationships were disrupted with the addition of strychnine, and recovered after washout; glycine current amplitude before and after washout varied similarly in the multi-omics context.

**Figure 3.**
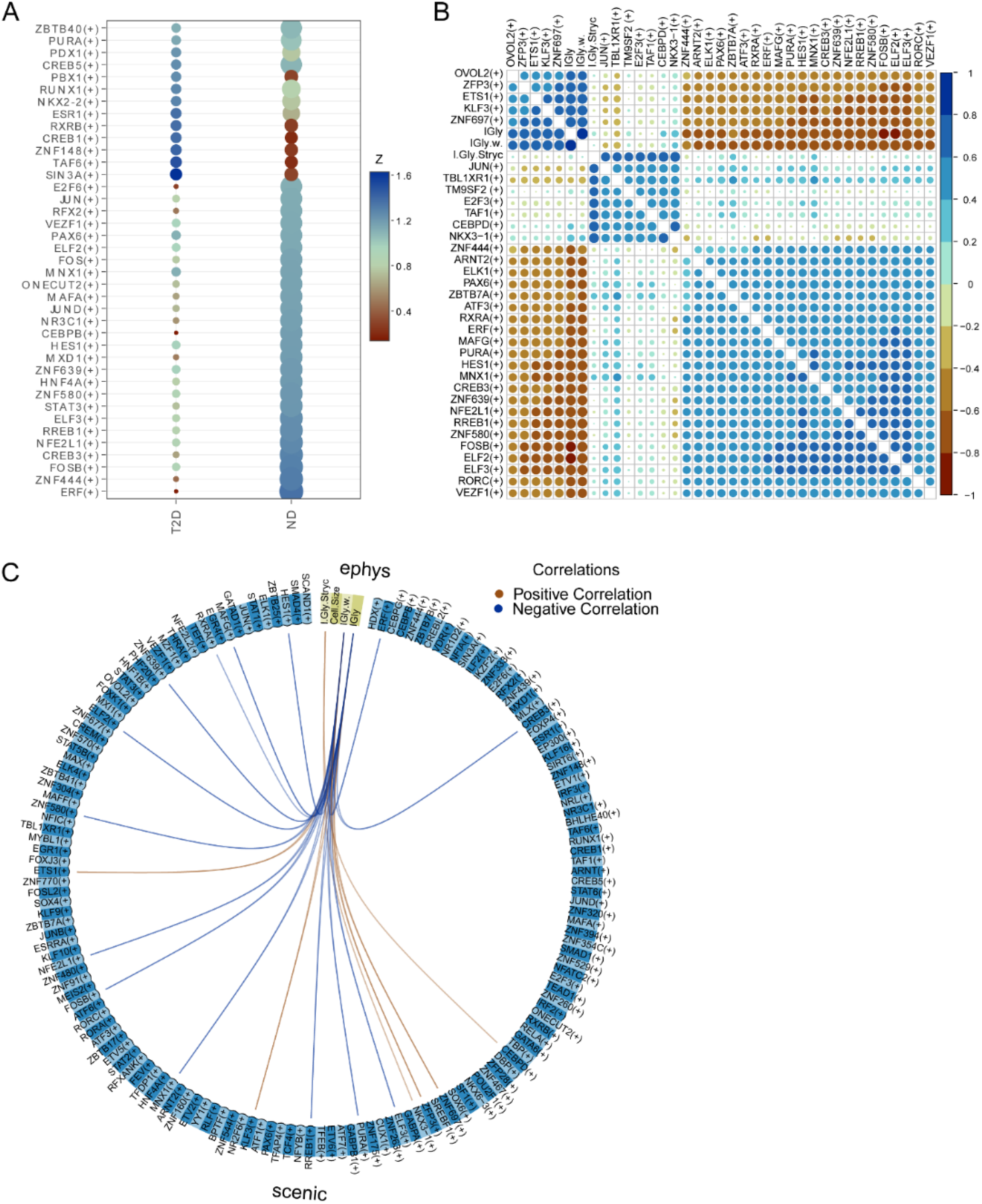
SCENIC and Multiomics Analysis. A: Dot plot showing regulon specificity score (RSS; size of dot) and Z score (colour of dot) of regulons found with SCENIC analysis in human β cells of donors with no diabetes (ND) and type 2 diabetes (T2D). Filtered by Z>1.105 and RSS>0.2. B: Similarity matrix dot plot showing correlation of features that maximize variation between human β cells of donors with ND and T2D from mixOmics analysis (both size of circle and colour indicate correlation). Correlation index scale is shown on the right. Filtered to only show features with abs(correlation) >0.6. C: Circos plot of mixOmics analysis, representing SCENIC regulons that are correlated with electrophysiology parameters. Features were selected to maximize variance between human β cells of donors with ND and T2D. Connections shown have an absolute correlation >0.7.

### Type 2 diabetes reduces β cell GlyR expression, but does not alter subunit splicing

We next investigated if the decreased GlyR currents in T2D are mediated by changes in the receptor expression or receptor composition, since the subunit expression could be shifted from functional to non-functional splice variants in T2D. For that, we quantified and compared the mRNA expression of all glycine receptor subunit splice variants in islets from donors with and without T2D, and after high glucose culture. The expression of the GlyR subunits α1 variant 1, 2, and 3, and α3 variant 1 are all significantly reduced in T2D islets (**Figure 4A**). The other variants (α1 variant 4, α3 variant 2, and β variants 1 and 3) showed similar levels of expression in nondiabetic and T2D islets. We also measured the expression of some splicing factors that could be responsible for GlyR splicing in islets, such as *NOVA1* and *PTBP2*, which regulate GlyR α2 subunit alternative splicing in neurons (48,49), and *PTBP1*, *NOVA2*, *RBFOX1*, *RBFOX2*, and *ELAVL4*, which are important splicing regulators of both neurons and β cells (50,51). No difference was seen in the RNA expression of the splicing factor genes *NOVA1*, *NOVA2*, *RBFOX1*, *RBFOX2*, *ELAVL4*, *PTBP1*, and *PTBP2* (**Figure 4B**). We confirm *GLRA1* downregulation in T2D (but not pre-diabetes) in an analysis using bulk RNAseq data at www.humanislets.com (52,53) (**Figure 5A**), together with an inverse correlation with donor HbA1c (p = 3.9×10^−5^, p-adj = 0.0275, coefficient = -10.2). The same is seen for the *GLRA3* gene, which encodes the α3 subunit (**Figure 5B**), also anticorrelated with donor HbA1c (p=0.024, p-adj = 0.361, coefficient = -3.96). Some of the transcripts correlated to glycine-evoked current amplitude have altered expression in islets from donors with T2D (**Figure 5C-I**).

**Figure 4.**
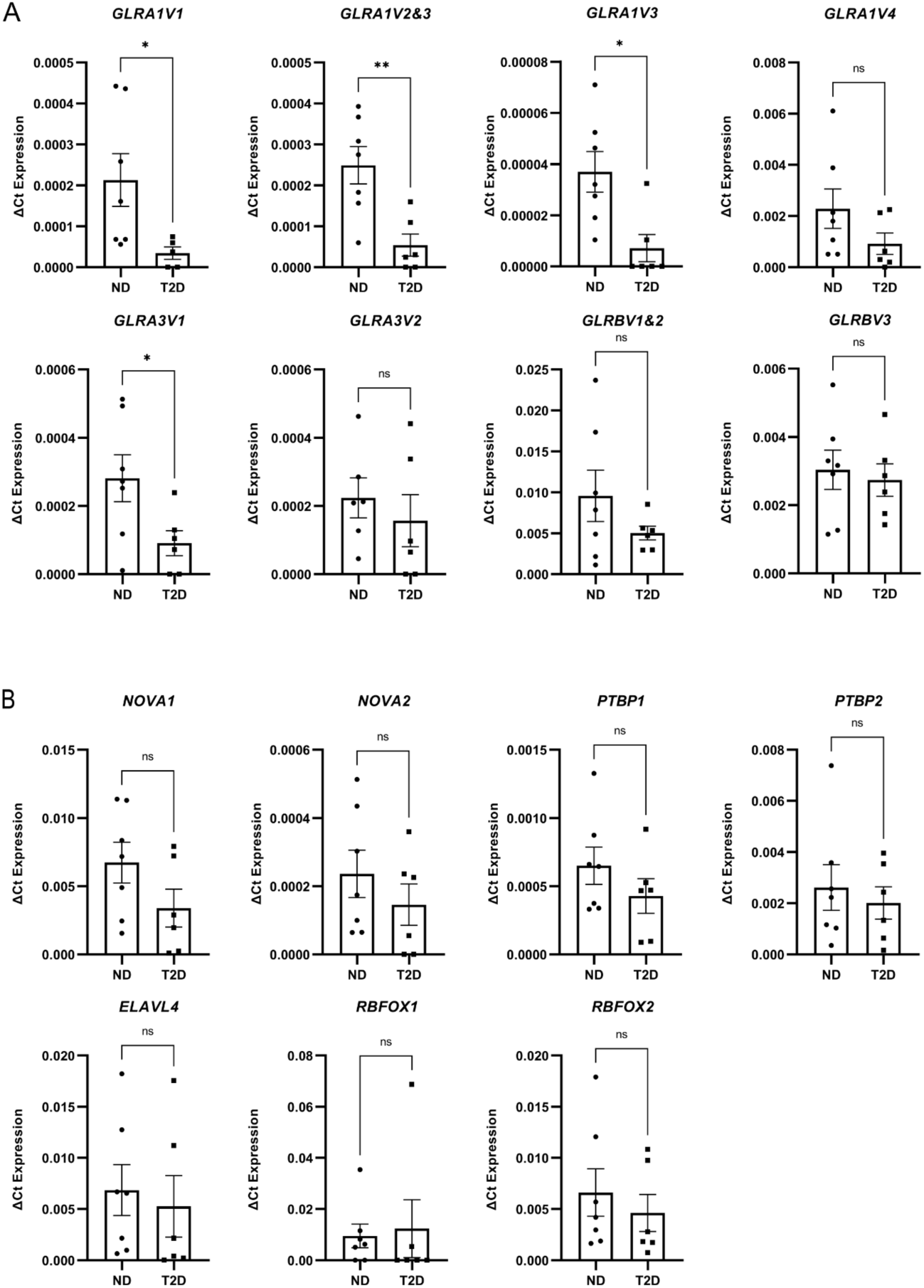
mRNA expression of GlyR subunit splice variants and splicing factors in human islets. A: Expression of GlyR subunits α1, α3 and β splice variants in islets of donors with (T2D, n = 6) and without T2D (ND, n = 7). Data was compared by unpaired t test or Mann Whitney test. B: mRNA expression of the splicing factors *NOVA1*, *NOVA2*, *RBFOX1*, *RBFOX2*, *ELAVL4*, *PTBP1* and *PTBP2* in human islets of donors with (n = 6) and without T2D (n = 7). Data is shown as ΔCt expression, calculated using the gene cyclophilin A (*PPIA*) as housekeeping. * P ≤ 0.05, ** P ≤ 0.01, and ns = non significant.

**Figure 5.**
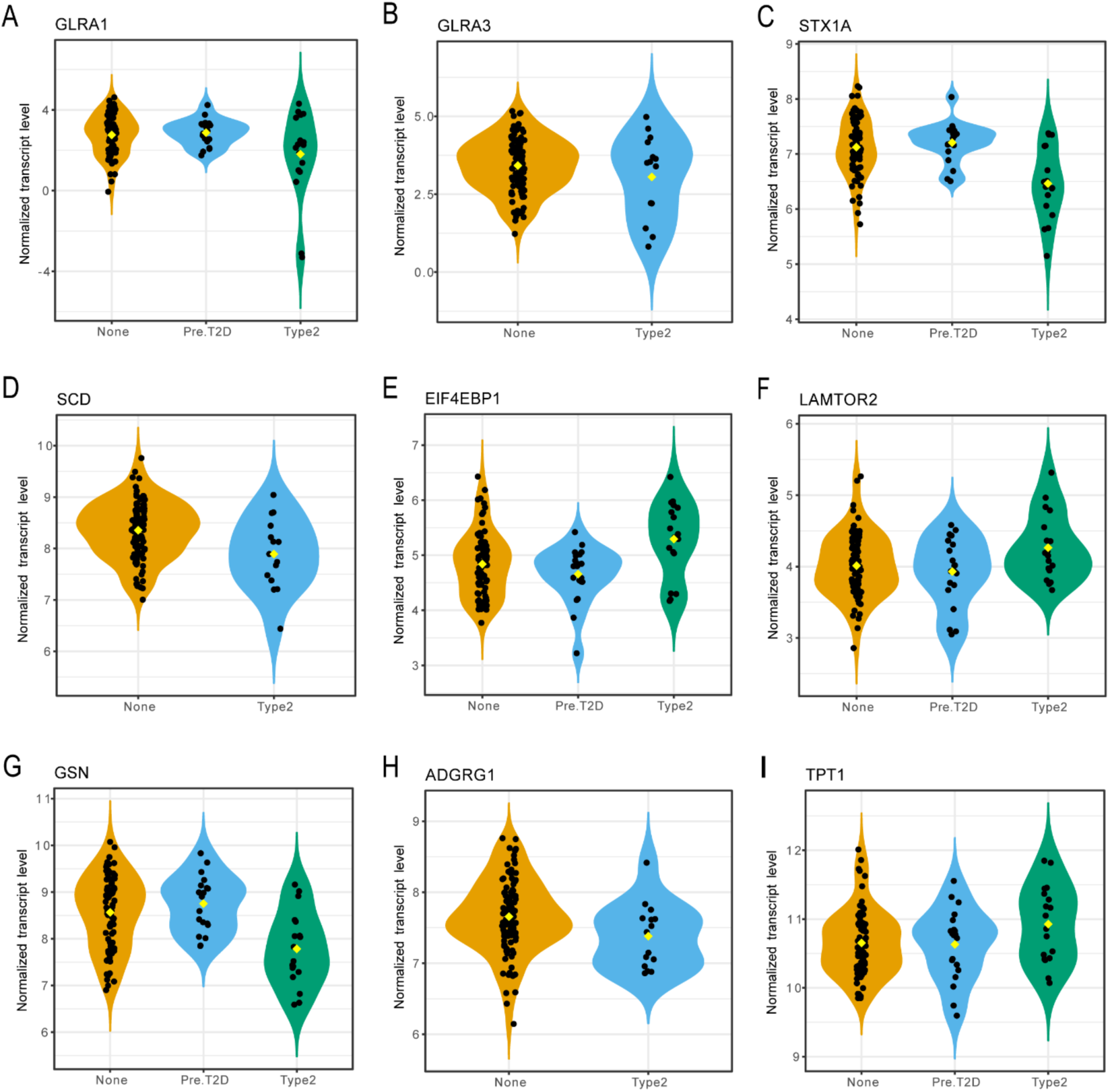
Bulk gene expression RNA-seq data from the www.humanislets.com website of transcripts correlated to glycine-induced current amplitude. Transcripts and GlyR subunits with altered expression between islets from donors with no diabetes (none), pre-type 2 diabetes (Pre.T2D), and/or type 2 diabetes (T2D). A: *GLRA1* (GlyR α1), p = 0.0003, p-adj = -0.0262, coefficient = -1.15. B: *GLRA3* (GlyR α3), p = 0.0266, p-adj = 0.188, coefficient = -0.525. C: *STX1A* (syntaxin 1A), p = 0.00000122, p-adj = 0.00219, coefficient = -0.714. D: *SCD* (stearoyl-CoA desaturase), p = 0.0101, p-adj = 0.107, coefficient = -0.412. E: *EIF4EBP1* (eukaryotic translation initiation factor 4E binding protein 1), p = 0.00227, p-adj = 0.0661, coefficient = 0.489. F: *LAMTOR2* (late endosomal/lysosomal adaptor, MAPK and MTOR activator 2), p = 0.0181, p-adj = 0.192, coefficient = 0.295. G: *GSN* (gelsolin), p = 0.000542, p-adj = 0.034, coefficient = - 0.708. H: *ADGRG1* (adhesion G protein-coupled receptor G1), p = 0.0337, p-adj = 0.214, coefficient = - 0.308. I: *TPT1* (tumor protein, translationally-controlled 1), p = 0.00438, p-adj = 0.0927, coefficient = 0.365.

### Hyperglycemia reduces β cell glycine-induced current

To investigate the effect of glycemia on β cell GlyRs, we first correlated glycine-induced current values of each β cell with its donor HbA1c (**Figure 6A,B**). We found that there is an inverse correlation between current amplitude and donor HbA1c, irrespective of diabetes status. No correlation was found between current amplitude and donor age and BMI, and there was no difference in the current between male and female donors (not shown). Next, to further verify if the GlyR dysfunction seen in type 2 diabetes could be caused by hyperglycemia, we measured glycine-evoked current after culturing human β cells from donors without diabetes in different glucose concentrations. The glycine-induced current was similar in cells cultured in 5.5 mM and 15 mM glucose for 24 hours (**Figure 6C**) but decreased after 48 hours of high glucose culture (**Figure 6D**). This was mirrored by a reduced expression of the GlyR subunits α1 variants 1, 3, and 4, while the β subunit showed no changes, and both α3 variants were increased (**Figure 6E**).

**Figure 6.**
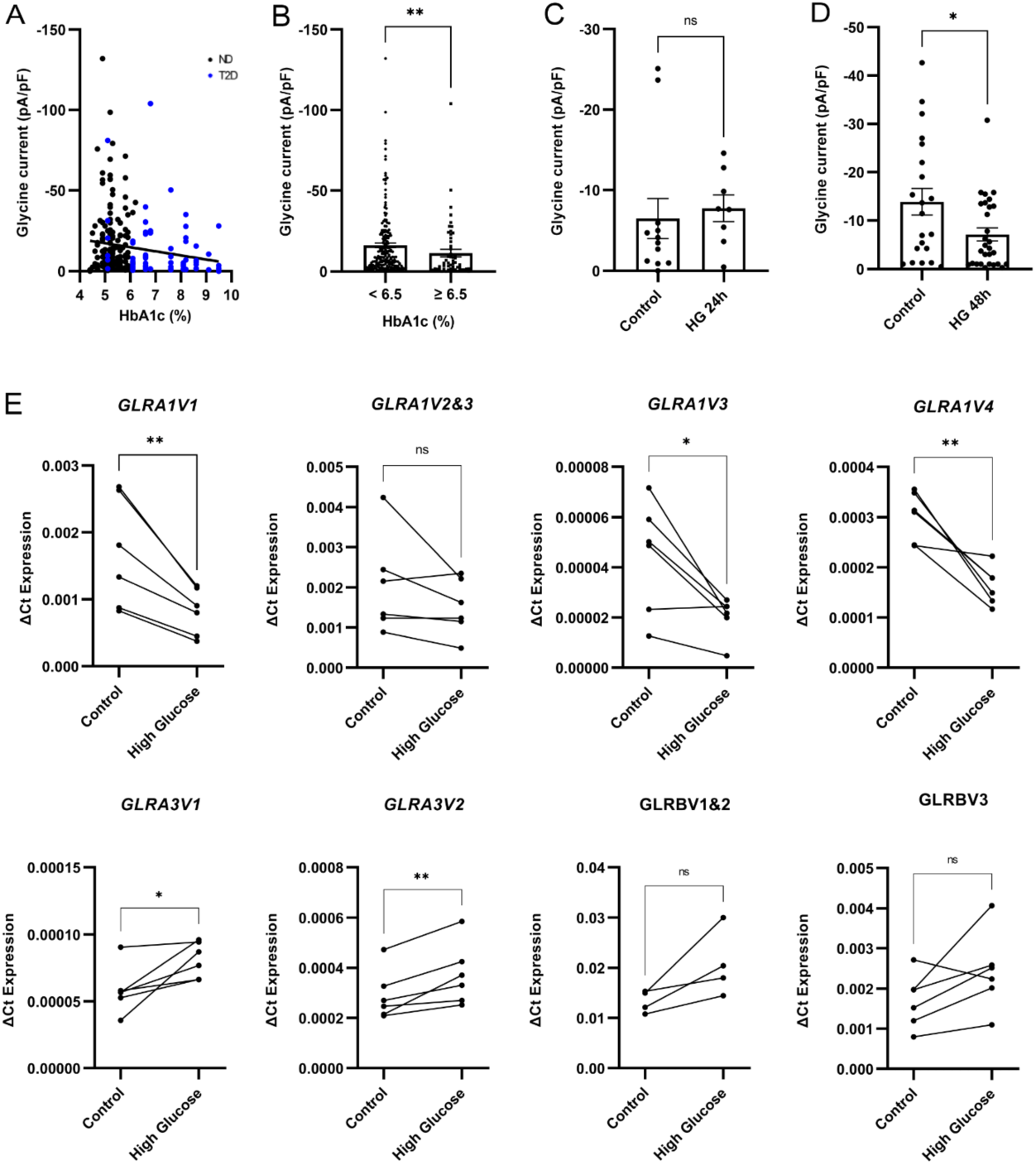
Glycine-induced currents and GlyR expression after high glucose culture. A: Correlation between donor HbA1c and glycine-induced current measured in each β cell. The correlation was calculated by simple linear regression (p = 0.0139, R squared = 0.02473) and Spearman correlation (P (two-tailed) = 0.0044, r = 0.1820, n = 244 cells). B: Glycine-induced current divided by donor HbA1c of < 6.5 (n=190 cells) and ≥ 6.5 (n=54 cells). Data was compared using the Mann-Whitney test, p = 0.0069. C: Currents from donors without diabetes cultured in 5.5 mM glucose (control, n = 12 cells from 7 donors) or 15 mM glucose for 24 hours (HG 24h, n = 8 cells from 5 donors). Data was compared using the Mann-Whitney test, p = 0.1813. D: Currents measured from donors without diabetes cultured for 48 hours in 5.5 mM glucose (n = 21 cells from 9 donors) or 15 mM glucose (HG 48h, n = 29 cells from 9 donors). Data was compared using the Mann-Whitney test, p = 0.0316. E: Expression of GlyR subunits α1, α3 and β mRNA splice variants in islets after 15 mM glucose culture for 48h (n = 6 donors, except for *GLRB* variants 1&2 (n=4 donors)). Data was compared by paired t test. Data is shown as ΔCt expression, calculated using the gene cyclophilin A as housekeeping. * P ≤ 0.05, ** P ≤ 0.01, ns = non significant.

## Discussion

Preserved paracrine and autocrine signaling, along with neurotransmitter input in the pancreatic islets is essential for the physiological control of insulin secretion, and its dysfunction can contribute to the pathophysiology of diabetes (54,55). Glycine was established to act in an autocrine positive feedback loop in human β cells (14), and we sought to investigate what causes its disruption in T2D and further characterize its action in β cells and the islets. We show that blocking GlyRs with strychnine reduced insulin secretion in human islets without the exogenous addition of glycine. This suggests that endogenous glycine from the islets, secreted from the α and/or β cells (14,15) has a tonic effect on glucose-stimulated insulin secretion. Even though some studies have shown that glycine ingestion raises insulin secretion (17,18) and that there is a negative correlation between plasma glycine levels and T2D (56), the exact glycine concentration in the pancreatic islets and whether it changes according to plasma glycine remains unknown. Efficient functioning of the glycine transporters expressed in β cells may indeed keep the inter-cellular glycine concentrations relatively low, as suggested to occur within the synapse (57,58).

We showed that glycine-evoked currents in human β cells are decreased in T2D, confirming previous findings (14). We then investigated if in T2D there is a change in which GlyR subunit mRNA splice variants are expressed, to elucidate if the GlyR dysfunction seen in this disease is due to the expression of less functional subunit splice variants. Our data shows that this is not the case, and that GlyR impairment in T2D is not due to a shift towards expression of less functional subunit variants, but more likely due to the overall decreased receptor gene expression. To test if hyperglycemia has a similar effect we cultured islet cells in 15 mM glucose, a supraphysiological concentration to simulate the chronic exposure to high glucose that happens in T2D, and measured GlyR gene expression and glycine-evoked currents. We discovered similar results in the gene expression of the GlyRs after high glucose culture as seen in T2D islets, where the α1 subunit was overall decreased while the β subunit remained unchanged. However, contrary to T2D islets, where the α3 subunit was downregulated, both α3 subunit variants were upregulated by glucose. This change in subunit likely does not result in up or down-regulated currents, since in recombinant systems heteromeric α1β and α3β receptors show similar channel conductance (59), and homomeric α1 and α3 receptors show similar maximal currents (60,61).

Consistent with our findings and the www.humanislets.com datasets (52,53), the *GLRA1* gene (which encodes the α1 subunit) was recently shown to be downregulated in islets from donors with T2D (36), in addition to having reduced expression after the exposure of islets to high glucose (33,34) and increased DNA methylation in the CpG sites (34). The exact intracellular mechanisms by which glucose causes these changes and decreases the receptor gene expression are still unclear. It has also been proposed that glycine receptors are altered in rat models of streptozotocin-induced diabetes, where GlyR expression was altered in retinal (62) and spinal cord neurons (63), and glycine neuronal signaling was impaired by pre-synaptic mechanisms (64), which suggests a glucose-dependent mechanism common to multiple cell types.

We found that after 48 hours of culture with high glucose, glycine-induced currents are reduced, which could be attributed to lower receptor expression. The β cells cultured in high glucose (15 mM) for 24 hours did not exhibit any difference in current amplitude. Given that the glycine receptors’ turnover time in the plasma membrane can have a half-life of 14, 24, or 48 hours (65–67), depending on the model used, it is possible that the 24-hour high glucose treatment was too short to reflect in a reduced number of receptors in the plasma membrane. Further establishing a relationship between glycine receptors and glucose is the inverse correlation found between donor HbA1c and glycine-induced current, suggesting that the main factor of T2D that is altering the GlyRs is hyperglycemia.

We found that glycine-induced current values in human β cells are highly variable and searched for molecules that could be involved in GlyR signaling modulation in β cells with the patch-seq technique. One inherent limitation of single-cell RNA sequencing is that it cannot capture 100% of a cell’s transcriptome, and especially transcripts with low expression levels tend to not be detected by the sequencing (68–70). Due to that fact, we did not detect the expression of the GlyR subunit genes themselves in single cells, which limited our ability to analyze if the cell-to-cell heterogeneity of glycine-evoked currents resulted from variability in the receptor expression levels of each cell. We did however find 92 transcripts positively or negatively correlated to glycine-induced current amplitude, both with previously known and unknown roles in the β cells. Some of the correlated genes found are altered by hyperglycemia or T2D, such as Translation initiation factor 4E-binding protein 1 (*EIF4EBP1*) (71,72), Polypyrimidine tract-binding protein 1 (*PTBP1*) (73,74), and Adhesion GPCR-G1/GPCR56 (*ADGRG1)* (75), and therefore could be involved in GlyR downregulation in T2D.

The SCENIC transcription factor analysis, coupled with the multi-omics integration, allows potential insights into the biology driving differences in glycine receptor activity. Regulons that were positively correlated with glycine receptor activity, indicating increased regulon activity with increased glycine receptor activity, included zinc finger proteins, such as OVOL2, ZNF697, and ZFP3. Together, these findings highlight how transcription factor activity and downstream targets may contribute to glycine receptor function in β cells, suggesting potential mechanisms of β cell dysfunction and stress in the context of T2D. We found that the activity of RREB1 is strongly linked to glycine receptor activity, with higher RREB1 activity correlated with lower glycine receptor activity. In this dataset, RREB1 is inferred to regulate *EIF4E*. Expression of *EIF4EBP1*, the binding protein that binds to and inhibits EIF4E, is strongly positively correlated with glycine receptor activity. Recently, RREB1 has been shown to influence beta cell function by transcriptionally regulating the expression of genes involved in β cell development and function (76). Thus, while glycine-evoked current is not significantly affected by silencing *EIF4EBP1* alone, upstream changes in RREB1 activity, or dysfunction, could be influencing glycine receptor impairment in T2D β cells. For example, regulator of G protein signaling protein 2 (*RGS2*), is known to have a role in controlling ion channel currents (77), and depletion of *RGS2* leads to insulin hypersecretion (78). Although the lower activity of RREB1 in β cells from donors with T2D would be predicted to lead to increased glycine current activity, the T2D β cells in the patch-seq data set did not have lower glycine receptor activity compared to the non-diabetic β cells; a larger dataset and targeted experiments to validate the roles of RREB1, EIF4EBP1, RGS2, and other transcripts would help clarify the regulatory network influencing glycine receptor activity in the context of T2D.

In summary, we demonstrate that glucose can modulate the GlyRs, by reducing their gene expression and consequently decreasing human β cell glycine-evoked current. Moreover, we define a previously unknown set of genes and regulons that are potentially related to GlyR function and could be involved in GlyR downregulation in T2D.

### CRediT Statement

Amanda Gomes: Conceptualization, Formal analysis, Investigation, Writing – Original Draft, Writing – Review & Editing, Visualization, Project administration. Patrick MacDonald: Supervision, Project administration, Conceptualization, Writing – Original Draft, Writing – Review & Editing, Funding acquisition, Resources. Aliya Spigelman: Investigation, Methodology. Kunimasa Suzuki: Investigation, Methodology. Theodore dos Santos: Formal analysis, Software. Jasmine Maghera: Methodology, Data curation. Cara Ellis: Formal analysis, Software, Investigation, Data curation, Writing – Review & Editing.

## Acknowledgements

The University of Alberta is situated on Treaty 6 territory, traditional lands of First Nations and Métis people. We thank the Human Organ Procurement and Exchange (HOPE) program and Trillium Gift of Life Network (TGLN) for their work in procuring human donor pancreas for research, and James Lyon (Alberta) for his efforts in human islet isolation. We especially thank the organ donors and their families for their kind gift in support of diabetes research. This work includes data and/or analyses from HumanIslets.com funded by the Canadian Institutes of Health Research, JDRF Canada, and Diabetes Canada (5-SRA-2021-1149-S-B/TG 179092) with data from islets isolated by the Alberta Diabetes Institute IsletCore with the support of the Human Organ Procurement and Exchange program, Trillium Gift of Life Network, BC Transplant, Quebec Transplant, and other Canadian organ procurement organizations with written informed donor consent as approved by the Human Research Ethics Board at the University of Alberta (Pro00013094).

This work was funded by a grant from the Canadian Institutes of Health Research to PEM (PS 186226). Some data used in this web tool includes patch-seq data, and single-cell RNA-seq used for cell type expression analysis, from the Human Pancreas Analysis Program (HPAP-RRID:SCR_016202) Database (https://hpap.pmacs.upenn.edu), a Human Islet Research Network (RRID:SCR_014393) consortium (UC4-DK-112217, U01-DK-123594, UC4-DK-112232, and U01-DK-123716).

ASG was supported in part by a studentship from Alberta-Helmholtz Diabetes Research School. JM was supported by the Vanier Canada Graduate Scholarship, the Alberta Innovates Graduate Studentship Scholarship, and the Sir Frederick Banting and Dr Charles Best Canada Graduate Scholarship. TdS was supported by the Alberta-Helmholtz Diabetes Research School, the Alberta Innovates Scholarship in Data-Enabled Innovation, and the Sir Fredrik Banting and Dr. Charles Best Canada Graduate Scholarship. PEM holds a Canada Research Chair in Islet Biology.

Finally, we thank Dr. Matthias Braun whose legacy inspired this work.

## References

1. Rorsman P, Ashcroft FM. Pancreatic β-Cell Electrical Activity and Insulin Secretion: Of Mice and Men. Physiol Rev. 2018 Jan 1;98(1):117–214.

2. Rorsman P, Braun M. Regulation of Insulin Secretion in Human Pancreatic Islets. Annu Rev Physiol. 2013 Feb 10;75(1):155–79.

3. Losada-Barragán M. Physiological effects of nutrients on insulin release by pancreatic beta cells. Mol Cell Biochem. 2021 Aug 12;476(8):3127–39.

4. Kolic J, Sun WG, Johnson JD, Guess N. Amino acid-stimulated insulin secretion: a path forward in type 2 diabetes. Amino Acids. 2023 Dec 15;55(12):1857–66.

5. Henquin JC. Paracrine and autocrine control of insulin secretion in human islets: evidence and pending questions. American Journal of Physiology-Endocrinology and Metabolism. 2021 Jan 1;320(1):E78–86.

6. Pan X, Tao S, Tong N. Potential Therapeutic Targeting Neurotransmitter Receptors in Diabetes. Front Endocrinol (Lausanne). 2022 May 20;13.

7. Hampton RF, Jimenez-Gonzalez M, Stanley SA. Unravelling innervation of pancreatic islets. Diabetologia. 2022 Jul 29;65(7):1069–84.

8. Huising MO. Paracrine regulation of insulin secretion. Diabetologia. 2020 Oct 7;63(10):2057–63.

9. Bennet H, Balhuizen A, Medina A, Dekker Nitert M, Ottosson Laakso E, Essén S, et al. Altered serotonin (5-HT) 1D and 2A receptor expression may contribute to defective insulin and glucagon secretion in human type 2 diabetes. Peptides (NY). 2015 Sep;71:113–20.

10. Taneera J, Jin Z, Jin Y, Muhammed SJ, Zhang E, Lang S, et al. γ-Aminobutyric acid (GABA) signalling in human pancreatic islets is altered in type 2 diabetes. Diabetologia. 2012 Jul 27;55(7):1985–94.

11. Zhu L, Rossi M, Doliba NM, Wess J. Beta-cell M3 muscarinic acetylcholine receptors as potential targets for novel antidiabetic drugs. Int Immunopharmacol. 2020 Apr;81:106267.

12. Marquard J, Otter S, Welters A, Stirban A, Fischer A, Eglinger J, et al. Characterization of pancreatic NMDA receptors as possible drug targets for diabetes treatment. Nat Med. 2015 Apr 16;21(4):363–72.

13. Gammelsaeter R, Frøyland M, Aragón C, Danbolt NC, Fortin D, Storm-Mathisen J, et al. Glycine, GABA and their transporters in pancreatic islets of Langerhans: evidence for a paracrine transmitter interplay. J Cell Sci. 2004 Aug 1;117(17):3749–58.

14. Yan-Do R, Duong E, Manning Fox JE, Dai X, Suzuki K, Khan S, et al. A Glycine-Insulin Autocrine Feedback Loop Enhances Insulin Secretion From Human β-Cells and Is Impaired in Type 2 Diabetes. Diabetes [Internet]. 2016 May 3;65(8):2311–21. Available from: 10.2337/db15-1272

15. Gammelsaeter R, Frøyland M, Aragón C, Danbolt NC, Fortin D, Storm-Mathisen J, et al. Glycine, GABA and their transporters in pancreatic islets of Langerhans: evidence for a paracrine transmitter interplay. J Cell Sci. 2004 Aug 1;117(17):3749–58.

16. Eulenburg V, Armsen W, Betz H, Gomeza J. Glycine transporters: essential regulators of neurotransmission. Trends Biochem Sci. 2005 Jun;30(6):325–33.

17. Gannon MC, Nuttall JA, Nuttall FQ. The metabolic response to ingested glycine. Am J Clin Nutr. 2002 Dec;76(6):1302–7.

18. González-Ortiz M, Medina-Santillán R, Martínez-Abundis E, Reynoso von Drateln C. Effect of Glycine on Insulin Secretion and Action in Healthy First-Degree Relatives of Type 2 Diabetes Mellitus Patients. Hormone and Metabolic Research. 2001 Jun;33(6):358–60.

19. Newgard CB, An J, Bain JR, Muehlbauer MJ, Stevens RD, Lien LF, et al. A Branched-Chain Amino Acid-Related Metabolic Signature that Differentiates Obese and Lean Humans and Contributes to Insulin Resistance. Cell Metab. 2009 Apr;9(4):311–26.

20. Takashina C, Tsujino I, Watanabe T, Sakaue S, Ikeda D, Yamada A, et al. Associations among the plasma amino acid profile, obesity, and glucose metabolism in Japanese adults with normal glucose tolerance. Nutr Metab (Lond). 2016 Dec 19;13(1):5.

21. Li X, Sun L, Zhang W, Li H, Wang S, Mu H, et al. Association of serum glycine levels with metabolic syndrome in an elderly Chinese population. Nutr Metab (Lond). 2018 Dec 17;15(1):89.

22. Glynn EL, Piner LW, Huffman KM, Slentz CA, Elliot-Penry L, AbouAssi H, et al. Impact of combined resistance and aerobic exercise training on branched-chain amino acid turnover, glycine metabolism and insulin sensitivity in overweight humans. Diabetologia. 2015 Oct 9;58(10):2324–35.

23. Merino J, Leong A, Liu CT, Porneala B, Walford GA, von Grotthuss M, et al. Metabolomics insights into early type 2 diabetes pathogenesis and detection in individuals with normal fasting glucose. Diabetologia. 2018 Jun 6;61(6):1315–24.

24. Gar C, Rottenkolber M, Prehn C, Adamski J, Seissler J, Lechner A. Serum and plasma amino acids as markers of prediabetes, insulin resistance, and incident diabetes. Crit Rev Clin Lab Sci. 2018 Jan 2;55(1):21–32.

25. Lynch JW. Native glycine receptor subtypes and their physiological roles. Neuropharmacology. 2009 Jan;56(1):303–9.

26. Raltschev C, Hetsch F, Winkelmann A, Meier JC, Semtner M. Electrophysiological Signature of Homomeric and Heteromeric Glycine Receptor Channels. Journal of Biological Chemistry. 2016 Aug;291(34):18030–40.

27. LI P, SLAUGHTER M. Glycine receptor subunit composition alters the action of GABA antagonists. Vis Neurosci. 2007 Jul 23;24(4):513–21.

28. Miller PS, Harvey RJ, Smart TG. Differential agonist sensitivity of glycine receptor *α* 2 subunit splice variants. Br J Pharmacol. 2004 Sep 17;143(1):19–26.

29. Dutertre S, Becker CM, Betz H. Inhibitory Glycine Receptors: An Update. Journal of Biological Chemistry. 2012 Nov;287(48):40216–23.

30. Braun M, Ramracheya R, Bengtsson M, Clark A, Walker JN, Johnson PR, et al. γ-Aminobutyric Acid (GABA) Is an Autocrine Excitatory Transmitter in Human Pancreatic β-Cells. Diabetes. 2010 Jul 1;59(7):1694–701.

31. Benninger RKP, Kravets V. The physiological role of β-cell heterogeneity in pancreatic islet function. Nat Rev Endocrinol. 2022 Jan 19;18(1):9–22.

32. Miranda MA, Macias-Velasco JF, Lawson HA. Pancreatic β-cell heterogeneity in health and diabetes: classes, sources, and subtypes. American Journal of Physiology-Endocrinology and Metabolism. 2021 Apr 1;320(4):E716–31.

33. Taneera J, Fadista J, Ahlqvist E, Atac D, Ottosson-Laakso E, Wollheim CB, et al. Identification of novel genes for glucose metabolism based upon expression pattern in human islets and effect on insulin secretion and glycemia. Hum Mol Genet. 2015 Apr 1;24(7):1945–55.

34. Hall E, Dekker Nitert M, Volkov P, Malmgren S, Mulder H, Bacos K, et al. The effects of high glucose exposure on global gene expression and DNA methylation in human pancreatic islets. Mol Cell Endocrinol. 2018 Sep;472:57–67.

35. Bacos K, Perfilyev A, Karagiannopoulos A, Cowan E, Ofori JK, Bertonnier-Brouty L, et al. Type 2 diabetes candidate genes, including PAX5, cause impaired insulin secretion in human pancreatic islets. Journal of Clinical Investigation. 2023 Feb 15;133(4).

36. Rönn T, Ofori JK, Perfilyev A, Hamilton A, Pircs K, Eichelmann F, et al. Genes with epigenetic alterations in human pancreatic islets impact mitochondrial function, insulin secretion, and type 2 diabetes. Nat Commun. 2023 Dec 12;14(1):8040.

37. Camunas-Soler J, Dai XQ, Hang Y, Bautista A, Lyon J, Suzuki K, et al. Patch-Seq Links Single-Cell Transcriptomes to Human Islet Dysfunction in Diabetes. Cell Metab. 2020 May;31(5):1017–1031.e4.

38. Lyon J, Manning Fox JE, Spigelman AF, Kim R, Smith N, O’Gorman D, et al. Research-Focused Isolation of Human Islets From Donors With and Without Diabetes at the Alberta Diabetes Institute IsletCore. Endocrinology [Internet]. 2016 Feb 1;157(2):560–9. Available from: 10.1210/en.2015-1562

39. Kin T, Shapiro J. Surgical aspects of human islet isolation. Islets [Internet]. 2010 Sep 1;2(5):265–73. Available from: 10.4161/isl.2.5.13019

40. Kaestner KH, Powers AC, Naji A, Consortium H, Atkinson MA. NIH Initiative to Improve Understanding of the Pancreas, Islet, and Autoimmunity in Type 1 Diabetes: The Human Pancreas Analysis Program (HPAP). Diabetes [Internet]. 2019 May 24;68(7):1394–402. Available from: 10.2337/db19-0058

41. Hagemann-Jensen M, Ziegenhain C, Chen P, Ramsköld D, Hendriks GJ, Larsson AJM, et al. Single-cell RNA counting at allele and isoform resolution using Smart-seq3. Nat Biotechnol. 2020 Jun;38(6):708–14.

42. Dobin A, Davis CA, Schlesinger F, Drenkow J, Zaleski C, Jha S, et al. STAR: ultrafast universal RNA-seq aligner. Bioinformatics. 2013 Jan 1;29(1):15–21.

43. Dai XQ, Camunas-Soler J, Briant LJB, dos Santos T, Spigelman AF, Walker EM, et al. Heterogenous impairment of α cell function in type 2 diabetes is linked to cell maturation state. Cell Metab. 2022 Feb;34(2):256–268.e5.

44. Van de Sande B, Flerin C, Davie K, De Waegeneer M, Hulselmans G, Aibar S, et al. A scalable SCENIC workflow for single-cell gene regulatory network analysis. Nat Protoc. 2020 Jul 19;15(7):2247–76.

45. Shrestha S, Erikson G, Lyon J, Spigelman AF, Bautista A, Manning Fox JE, et al. Aging compromises human islet beta cell function and identity by decreasing transcription factor activity and inducing ER stress. Sci Adv. 2022 Oct 7;8(40).

46. Rohart F, Gautier B, Singh A, Lê Cao KA. mixOmics: An R package for ‘omics feature selection and multiple data integration. PLoS Comput Biol. 2017 Nov 3;13(11):e1005752.

47. Guo S, Dai C, Guo M, Taylor B, Harmon JS, Sander M, et al. Inactivation of specific β cell transcription factors in type 2 diabetes. Journal of Clinical Investigation. 2013 Aug 1;123(8):3305–16.

48. Kumar D V, Nighorn A, St. John PA. Role of nova-1 in regulating α2N, a novel glycine receptor splice variant, in developing spinal cord neurons. J Neurobiol [Internet]. 2002 Aug 1;52(2):156–65. Available from: 10.1002/neu.10072

49. Polydorides AD, Okano HJ, Yang YYL, Stefani G, Darnell RB. A brain-enriched polypyrimidine tract-binding protein antagonizes the ability of Nova to regulate neuron-specific alternative splicing. Proceedings of the National Academy of Sciences [Internet]. 2000 Jun 6;97(12):6350–5. Available from: 10.1073/pnas.110128397

50. Dlamini Z, Mokoena F, Hull R. Abnormalities in alternative splicing in diabetes: therapeutic targets. J Mol Endocrinol [Internet]. 2017;59(2):R93–107. Available from: https://jme.bioscientifica.com/view/journals/jme/59/2/JME-17-0049.xml

51. Moss ND, Wells KL, Theis A, Kim YK, Spigelman AF, Liu X, et al. Modulation of insulin secretion by RBFOX2-mediated alternative splicing. Nat Commun [Internet]. 2023;14(1):7732. Available from: 10.1038/s41467-023-43605-4

52. Ewald JD, Lu Y, Ellis CE, Worton J, Kolic J, Sasaki S, et al. HumanIslets: An integrated platform for human islet data access and analysis. bioRxiv [Internet]. 2024 Jan 1;2024.06.19.599613. Available from: http://biorxiv.org/content/early/2024/06/22/2024.06.19.599613.abstract

53. Kolic J, Sun WG, Cen HH, Ewald JD, Rogalski JC, Sasaki S, et al. Proteomic predictors of individualized nutrient-specific insulin secretion in health and disease. Cell Metab. 2024 Jul;36(7):1619–1633.e5.

54. Huising MO. Paracrine regulation of insulin secretion. Diabetologia. 2020 Oct 7;63(10):2057–63.

55. Hartig SM, Cox AR. Paracrine signaling in islet function and survival. J Mol Med. 2020 Apr 17;98(4):451–67.

56. Yan-Do R, MacDonald PE. Impaired “glycine”-mia in type 2 diabetes and potential mechanisms contributing to glucose homeostasis. Vol. 158, Endocrinology. Endocrine Society; 2017. p. 1064–73.

57. Beato M. The time course of transmitter at glycinergic synapses onto motoneurons. J Neurosci. 2008 Jul 16;28(29):7412–25.

58. Harsing LG, Matyus P. Mechanisms of glycine release, which build up synaptic and extrasynaptic glycine levels: the role of synaptic and non-synaptic glycine transporters. Brain Res Bull. 2013 Apr;93:110–9.

59. Bormann J, Rundström N, Betz H, Langosch D. Residues within transmembrane segment M2 determine chloride conductance of glycine receptor homo- and hetero-oligomers. EMBO J. 1993 Oct;12(10):3729–37.

60. Miller PS, Beato M, Harvey RJ, Smart TG. Molecular determinants of glycine receptor αβ subunit sensitivities to Zn ^2+^ -mediated inhibition. J Physiol. 2005 Aug 27;566(3):657–70.

61. Chen X, Webb TI, Lynch JW. The M4 transmembrane segment contributes to agonist efficacy differences between α1 and α3 glycine receptors. Mol Membr Biol. 2009 Jan 19;26(5–7):321–32.

62. Morales-Calixto E, Velázquez-Flores MÁ, Sánchez-Chávez G, Ruiz Esparza-Garrido R, Salceda R. Glycine receptor is differentially expressed in the rat retina at early stages of streptozotocin-induced diabetes. Neurosci Lett. 2019 Nov;712:134506.

63. Velázquez-Flores MÁ, Sánchez-Chávez G, Morales-Lázaro SL, Ruiz Esparza-Garrido R, Canizales-Ontiveros A, Salceda R. Streptozotocin-Induced Diabetic Rats Showed a Differential Glycine Receptor Expression in the Spinal Cord: A GlyR Role in Diabetic Neuropathy. Neurochem Res. 2024 Mar 29;49(3):684–91.

64. Chiu YC, Liao WT, Liu CK, Wu CH, Lin CR. Reduction of spinal glycine receptor-mediated miniature inhibitory postsynaptic currents in streptozotocin-induced diabetic neuropathic pain. Neurosci Lett. 2016 Jan;611:88–93.

65. Rasmussen H, Rasmussen T, Triller A, Vannier C. Strychnine-Blocked Glycine Receptor Is Removed from Synapses by a Shift in Insertion/Degradation Equilibrium. Molecular and Cellular Neuroscience. 2002 Feb;19(2):201–15.

66. Villmann C, Oertel J, Melzer N, Becker C. Recessive hyperekplexia mutations of the glycine receptor α1 subunit affect cell surface integration and stability. J Neurochem. 2009 Nov 12;111(3):837–47.

67. Hoch W, Betz H, Becker CM. Primary cultures of mouse spinal cord express the neonatal isoform of the inhibitory glycine receptor. Neuron. 1989 Sep;3(3):339–48.

68. Ngara M, Wierup N. Lessons from single-cell RNA sequencing of human islets. Diabetologia. 2022 Aug 28;65(8):1241–50.

69. Wang YJ, Kaestner KH. Single-Cell RNA-Seq of the Pancreatic Islets––a Promise Not yet Fulfilled? Cell Metab. 2019 Mar;29(3):539–44.

70. Mawla AM, Huising MO. Navigating the Depths and Avoiding the Shallows of Pancreatic Islet Cell Transcriptomes. Diabetes. 2019 Jul 1;68(7):1380–93.

71. Yuan T, Rafizadeh S, Gorrepati KDD, Lupse B, Oberholzer J, Maedler K, et al. Reciprocal regulation of mTOR complexes in pancreatic islets from humans with type 2 diabetes. Diabetologia. 2017 Apr 21;60(4):668–78.

72. Yamaguchi S, Ishihara H, Yamada T, Tamura A, Usui M, Tominaga R, et al. ATF4-Mediated Induction of 4E-BP1 Contributes to Pancreatic β Cell Survival under Endoplasmic Reticulum Stress. Cell Metab. 2008 Mar;7(3):269–76.

73. Fred RG, Bang-Berthelsen CH, Mandrup-Poulsen T, Grunnet LG, Welsh N. High Glucose Suppresses Human Islet Insulin Biosynthesis by Inducing miR-133a Leading to Decreased Polypyrimidine Tract Binding Protein-Expression. PLoS One. 2010 May 26;5(5):e10843.

74. Jeong DE, Heo S, Han JH, Lee EY, Kulkarni RN, Kim W. Glucose Controls the Expression of Polypyrimidine Tract-Binding Protein 1 via the Insulin Receptor Signaling Pathway in Pancreatic β Cells. Mol Cells. 2018 Oct 31;41(10):909–16.

75. Dunér P, Al-Amily IM, Soni A, Asplund O, Safi F, Storm P, et al. Adhesion G Protein-Coupled Receptor G1 (ADGRG1/GPR56) and Pancreatic β-Cell Function. J Clin Endocrinol Metab. 2016 Dec;101(12):4637–45.

76. Mattis KK, Krentz NAJ, Metzendorf C, Abaitua F, Spigelman AF, Sun H, et al. Loss of RREB1 in pancreatic beta cells reduces cellular insulin content and affects endocrine cell gene expression. Diabetologia. 2023 Apr 12;66(4):674–94.

77. Nguyen CH, Ming H, Zhao P, Hugendubler L, Gros R, Kimball SR, et al. Translational control by RGS2. Journal of Cell Biology. 2009 Sep 7;186(5):755–65.

78. Dong H, Zhang Y, Wang J, Kim DS, Wu H, Sjögren B, et al. Regulator of G protein signaling 2 is a key regulator of pancreatic β-cell mass and function. Cell Death Dis. 2017 May 25;8(5):e2821–e2821.

